# Anaerobic metabolism of Foraminifera thriving below the seafloor

**DOI:** 10.1101/2020.03.26.009324

**Authors:** William D. Orsi, Raphaël Morard, Aurele Vuillemin, Michael Eitel, Gert Wörheide, Jana Milucka, Michal Kucera

## Abstract

Foraminifera are single-celled eukaryotes (protists) of large ecological importance, as well as environmental and paleoenvironmental indicators and biostratigraphic tools. In addition, they are capable of surviving in anoxic marine environments where they represent a major component of the benthic community. However, the cellular adaptations of Foraminifera to the anoxic environment remain poorly constrained. We sampled an oxic-anoxic transition zone in marine sediments from the Namibian shelf, where the genera *Bolivina* and *Stainforthia* dominated the Foraminifera community, and use metatranscriptomics to characterize Foraminifera metabolism across the different geochemical conditions. The relative abundance of Foraminifera gene expression in anoxic sediment depths increased an order of magnitude, which was confirmed in a ten-day incubation experiment where the development of anoxia coincided with a 27-fold increase in the relative abundance of Foraminifera protein encoding transcripts. This indicates that many Foraminifera were not only surviving, but thriving under the anoxic conditions. The anaerobic energy metabolism of these active Foraminifera was characterized by fermentation of sugars and amino acids, dissimilatory nitrate reduction, fumarate reduction, and dephosphorylation of creatine phosphate. This was co-expressed alongside genes involved in production of reticulopodia, phagocytosis, calcification, and clathrin-mediated-endocytosis (CME). Thus, Foraminifera may use CME under anoxic conditions to utilize dissolved organic matter as a carbon and energy source, in addition to ingestion of prey cells via phagocytosis. These mechanisms help explain how some Foraminifera can thrive under anoxia, which would help to explain their ecological success documented in the fossil record since the Cambrian period more than 500 million years ago.

## Introduction

Foraminifera are one of the most ubiquitous free-living marine eukaryotes on Earth and have been documented in the fossil record since the Cambrian period (*1*), surviving all mass extinction events involving extensive ocean anoxia (*2*). Benthic foraminifera inhabit marine sediments (*3*), where they can represent up to 50% of the sediment biomass in shallow depths of the seabed (*4*) and play a significant role in the benthic carbon and nitrogen cycles (*5*). Foraminifera are known to be resistant to oxygen depletion and may persist in the benthic community even under the development of anoxic and sulfidic conditions (*6-8*). A key to their survival in the absence of oxygen is their ability to perform complete denitrification (*9*), which appears to be a shared trait among many clades that likely evolved early in the evolutionary history of the group (*10*). A better understanding of anaerobic metabolism in Foraminifera under anoxic conditions could illuminate their ecological role in the benthos (*11*) and explain the ecological success of Foraminifera throughout the Phanerozoic, across multiple mass extinction events and associated widespread ocean anoxia (*2*).

To this end, we applied metatranscriptomics to study the active gene expression of anaerobic benthic Foraminifera in anoxic Namibian shelf sediments, and reconstruct their active biochemical pathways in situ. Our transcriptomic analysis showed the anaerobic pathways of ATP production, and revealed the biosynthetic processes that consume ATP. Our data indicate that Foraminifera are not only surviving under anoxic conditions, but that their activity is stimulated by anoxia. Analysis further shows the anaerobic mechanisms of ATP production which benthic Foraminifera employ to produce sufficient energy to power a multitude of energetically expensive cellular processes in the absence of oxygen. Transcriptional activity could be stimulated by the development of anoxic conditions during a ten day incubation indicating that many benthic Foraminifera are not only surviving, but appear to thrive under anoxic conditions.

## Results

A total of 14 sediment depth horizons were analyzed from a 28 cm long sediment core sectioned every 2 cm, which was retrieved from 125 m water depth on the continental shelf off Namibia (*12*). The core was sampled (sliced) immediately and stored at -20 C within 30 minutes after collection for metatranscriptomics and quantitative community composition estimates via microscopy. The pore water chemical analysis indicated that nitrate and nitrite were consumed quickly at the sediment surface followed by an increased accumulation of ammonium and sulfide with depth (Fig 1). Intact Foraminifera cells containing cytoplasm observed with light microscopy decreased in abundance with increasing depth, but were still present in the deepest part of the core indicating that these Foraminifera cells were living under anoxic conditions (Fig. 1). However, burrowing polychaete worms were observed throughout the core indicating the potential for downward vertical transport of oxidized porewater (e.g., containing O_2_, NO_3_^-^) via bioirrigation processes. Throughout the entire core sequence, 95% of the Foraminifera community at all depths was represented by the genera *Bolivina* and *Stainforthia*. We observed a bimodal distribution of the foraminifera absolute abundance with the maximum density at the oxic-anoxic transition at the surface layer of with ∼ 260 benthic foraminifera individuals per gram of sediment, followed by a steep decrease until 12-14 centimeters below sea floor (cmbsf) with 30 individuals per gram of sediment followed by an increase to 80 individuals per gram of sediment at 20-22 cmbsf, coinciding with nitrate-sulfide transition zone (Fig 1).

**Figure 1.**
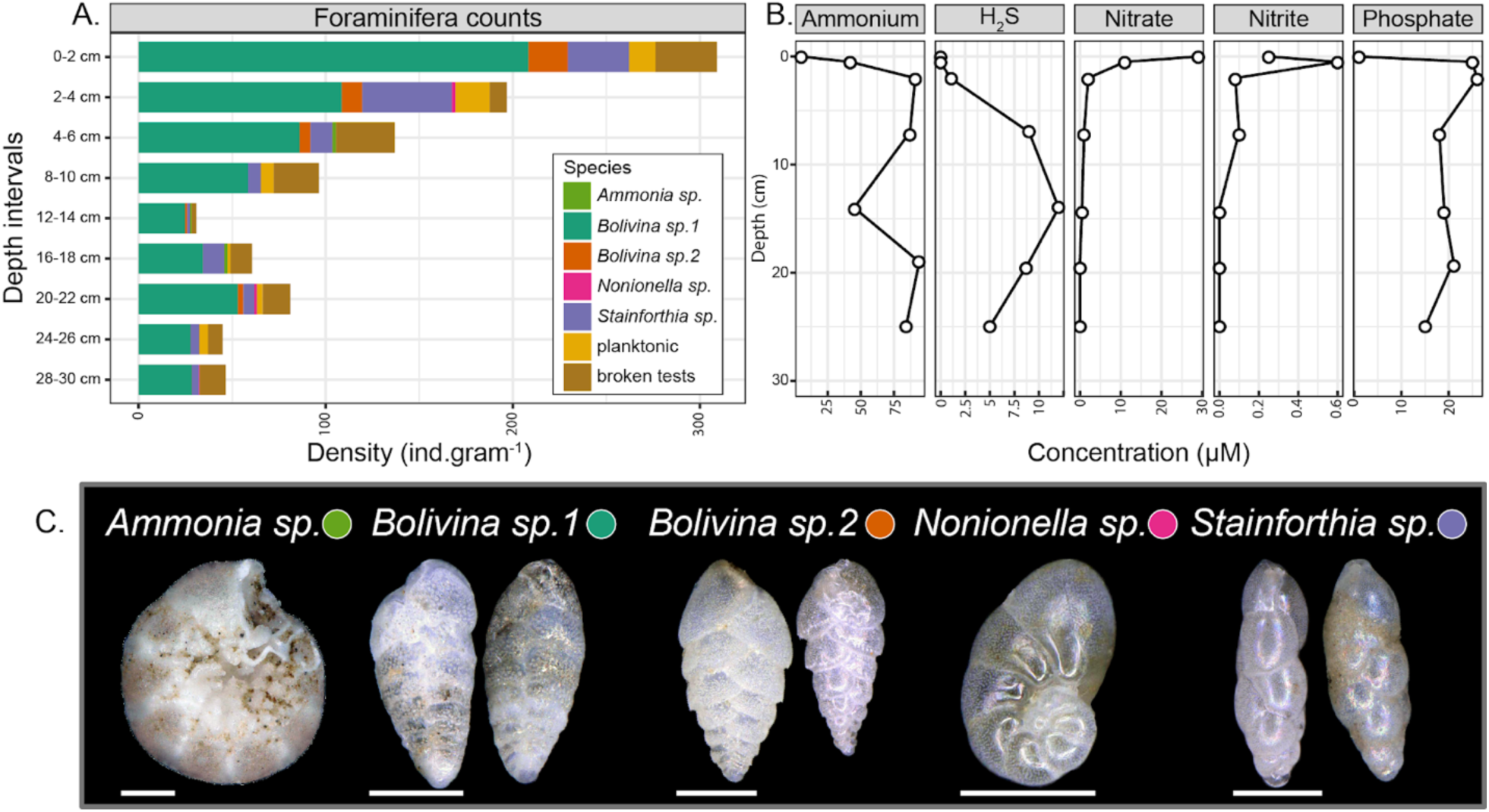
Census count of cytoplasm-containing foraminifera tests and corresponding geochemical profiles in anoxic Namibian sediment. (A) Density of the foraminifera species in the nine intervals processed compared against (B) the changing redox profile of in sediment pore water, note the accumulation of hydrogen sulphide with depth below 6 cm. (C) Representative specimens of the species enumerated, brownish-green color indicates the presence of cytoplasm. Scale bar 100 µm.

Metatranscriptomes were sequenced to a depth of on average 6 (+/- 5) million reads per sample (Table S1). Analyses of the metatranscriptomes showed that the Foraminifera increased their gene expression significantly under anoxic conditions, and they exhibited levels of gene expression far greater than all other groups of protists identified in the transcriptomes (Fig 2). The absolute level of gene expression by the Foraminifera increased with depth, because the total number of unique expressed protein encoding open reading frames (ORFs) assigned to Foraminifera increased (Fig 2b). An higher number of absolute unique ORFs expressed by Foraminifera cannot be explained by a reduction in gene expression from other groups. Clearly, some of the Foraminifera that were observed with intact cytoplasm in the deeper part of the core (Fig 1) increase their gene expression under anoxic conditions (Fig 2b, c).

**Figure 2.**
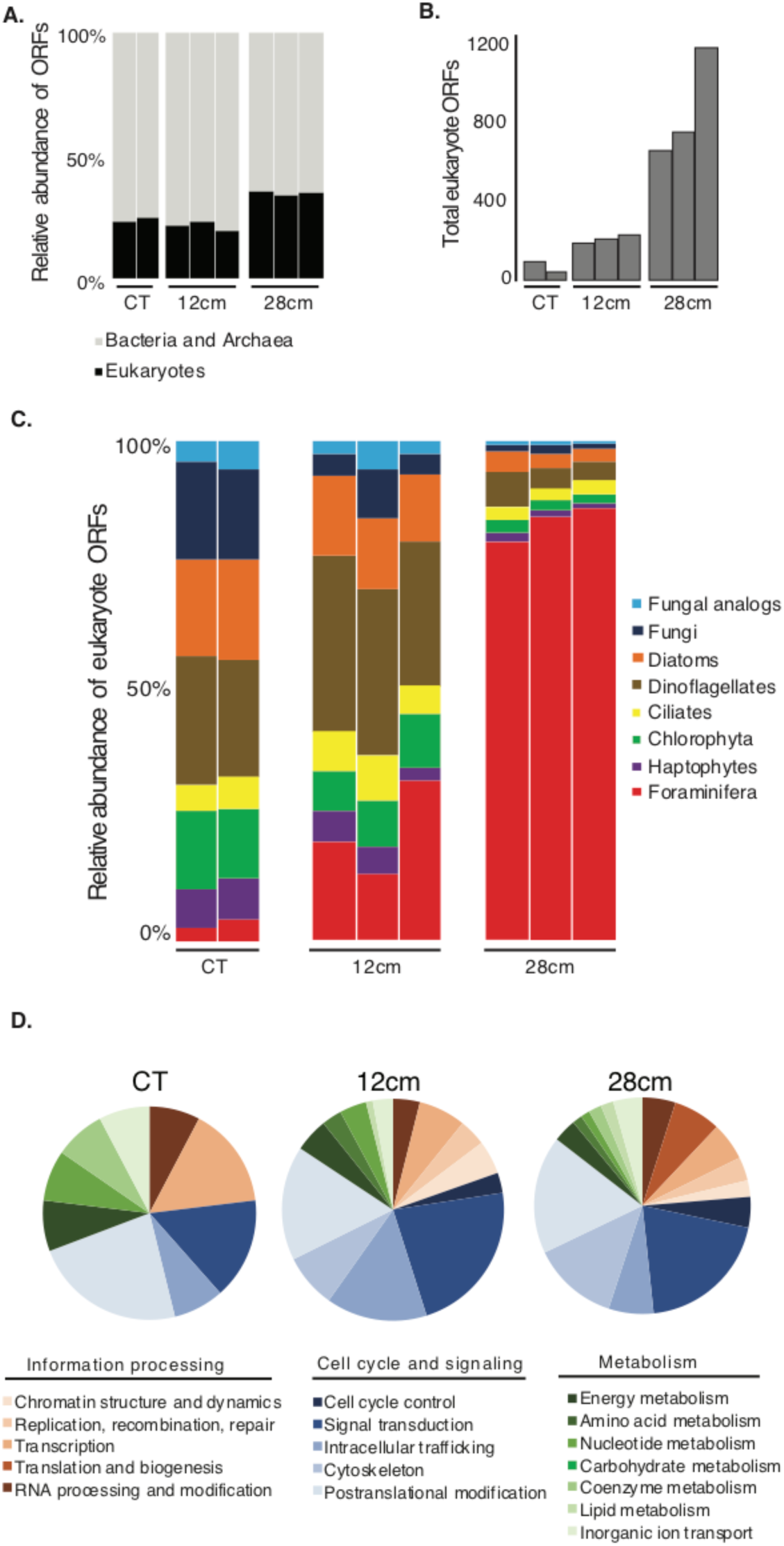
Foraminifera exhibit high levels of gene expression under anoxia. (A) The relative abundance of total expressed ORFs per sample that were assigned to prokaryotes (Bacteria and Archaea) and eukaryotes (including Foraminifera). Multiple histograms per depth represent biological replicates. (B) The total number of ORFs that were assigned to eukaryotes per sample. Multiple histograms per depth represent biological replicates. (C) The relative abundance of expressed ORFs from different protist Phyla (from panel B), note the dominance of Foraminifera gene expression in the deepest, most anoxic sample at 28 cm. (D) The relative abundance of functional eukaryotic gene (KOG) families in the three sediment zones that were assigned to expressed Foraminifera ORFs. Pie charts represent average values from the biological replicates shown in panels A-C. CT: core top sample.

Phylogenetic analyses of two Foraminifera 18S rDNA sequences recovered from the metatranscriptomes had closest affiliation to previously reported *Stainforthia* and *Bolivina* 18S rDNA sequences, also recovered from anoxic Namibian sediments (Fig. 3). *Stainforthia* and *Bolivina* tests containing cytoplasm were also observed in the core, their relative abundance gradually increased with depth, and *Bolivina* was the most abundant genus observed (Fig 1). Successful detection of its expressed 18S rRNA confirms that our metatranscriptomic approach captured the activity of this numerically dominant group. This is also reflected by the read mapping statistics (Figure S2), which support the ratios observed based on counts of cytoplasm containing tests with the *Bolivina* sp. 18S rRNA fragment showing a maximum read coverage of 312x and an average coverage of 125x. In contrast, the 18S rRNA from the comparatively less abundant cytoplasm containing tests from *Stainforthia* sp. (Fig 1) had lower maximum and mean coverages 135x and 34x, respectively. In contrast to 18S rRNA sequences, metatranscriptomic ORFs had the highest similarity to previously sequenced genomes and transcriptomes of *Ammonia, Elphidium, Rosalina*, and *Globobulimina* cells (Fig S1), the very few previously sequenced transcriptomes derived from Foraminifera (*10, 13, 14*). We could not find publicly available genome or transcriptome data from *Stainforthia* or *Bolivina* to include in our database for annotating the metatranscriptome data. Thus given that we could only detect 18S rRNA from *Stainforthia* and *Bolivina* (Fig 3) in the metatranscriptomes (and none from *Ammonia, Elphidium, Rosalina*, and *Globobulimina)*, we assume that most of the ORFs with highest similarity to Foraminifera are likely derived from the numerically dominant *Stainforthia* and *Bolivina* cells observed in the core (Fig 1), but have top hits to other Foraminifera (e.g., *Ammonia, Elphidium, Rosalina*, and *Globobulimina*: Fig S1) since *Stainforthia* and *Bolivina* transcriptomes are missing in our database. We then proceeded to analyze these Foraminifera-derived ORFs in the metatranscriptomes to gain insights into possibly anaerobic biochemical pathways and physiologies, after annotating all of the Foraminifera-derived ORFs against the clusters of Eukaryotic Orthologous Genes (KOGs) database (*15*).

**Figure 3.**
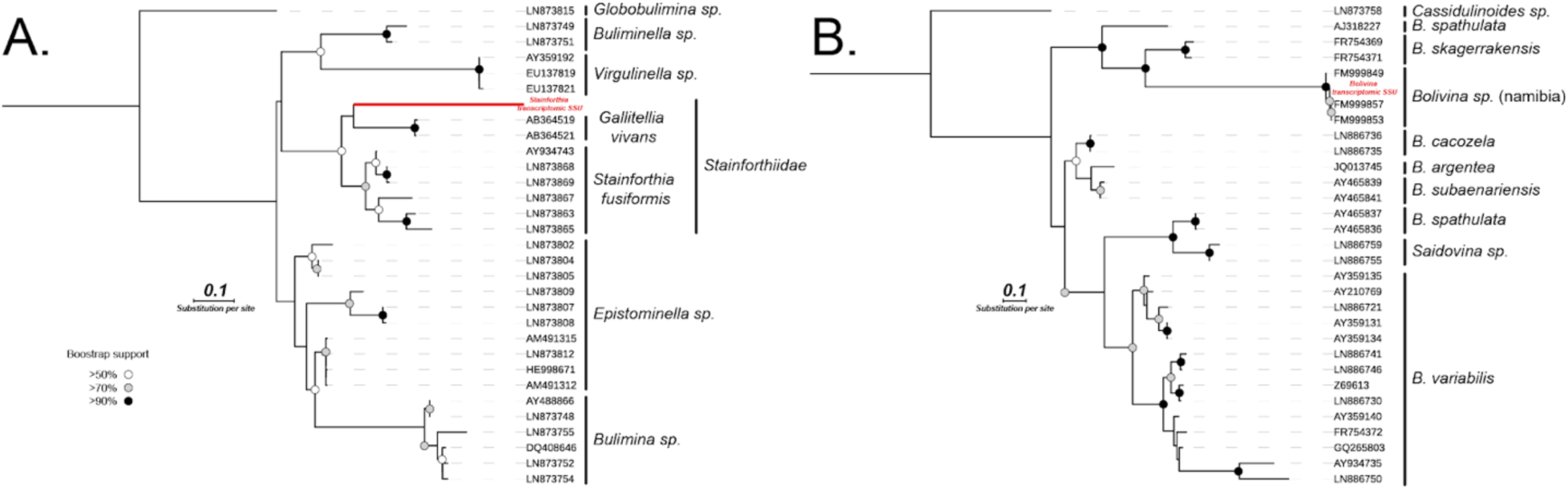
Phylogenetic analysis of Foraminifera affiliated 18S rDNA sequences recovered from the metatranscriptome, that are affiliated to the (A) Stainforthiidae family and (B) *Bolivina* genus. The sequence affiliated to the Stainforthiidae family clearly cluster with the only two representative genus of the family, *Stainforthia* and *Gallietellia* but the position of the metatranscriptomic 18S rDNA sequence is not clearly resolved, but intact test of *Stainforthia* were observed in the sample (See Fig. 1). The metatranscriptomic 18S rDNA sequence related to *Bolivina* is nearly identical to reference sequences deposited on NCBI and that were generated from *Bolivina* specimens collected in Namibia in previous studies. Furthermore, *Bolivina* specimens dominated the morphological assemblages within the core (Fig. 1). The *Bolivina* and *Stainforthia* 18S rDNA contigs were generated by semi-automated greedy extension of 18S rDNA OTUs with trimmed metatranscriptomic paired-end reads (See methods).

Expression of foraminiferal KOGs showed that at all depths the transcriptional activity was dominated by genes involved in cell cycle and cell signaling processes, namely cell cycle control, signal transduction, intracellular trafficking, cytoskeleton, and posttranslational modification (Fig 2). The expression of genes involved in translation and biogenesis was detected only in the deepest, anoxic sample indicating an increase in growth and biosynthesis in Foraminifera cells. There was also a general trend of decreasing energy production and conversion (COG category C) with depth, together with an increasing expression of genes involved in signal transduction under anoxic conditions (Fig 2). The gene expression from Foraminifera was significantly different between the anoxic depth at 28 cmbsf, and the other shallower depths (Fig 4a: ANOSIM, *P* < 0.01).

**Figure 4.**
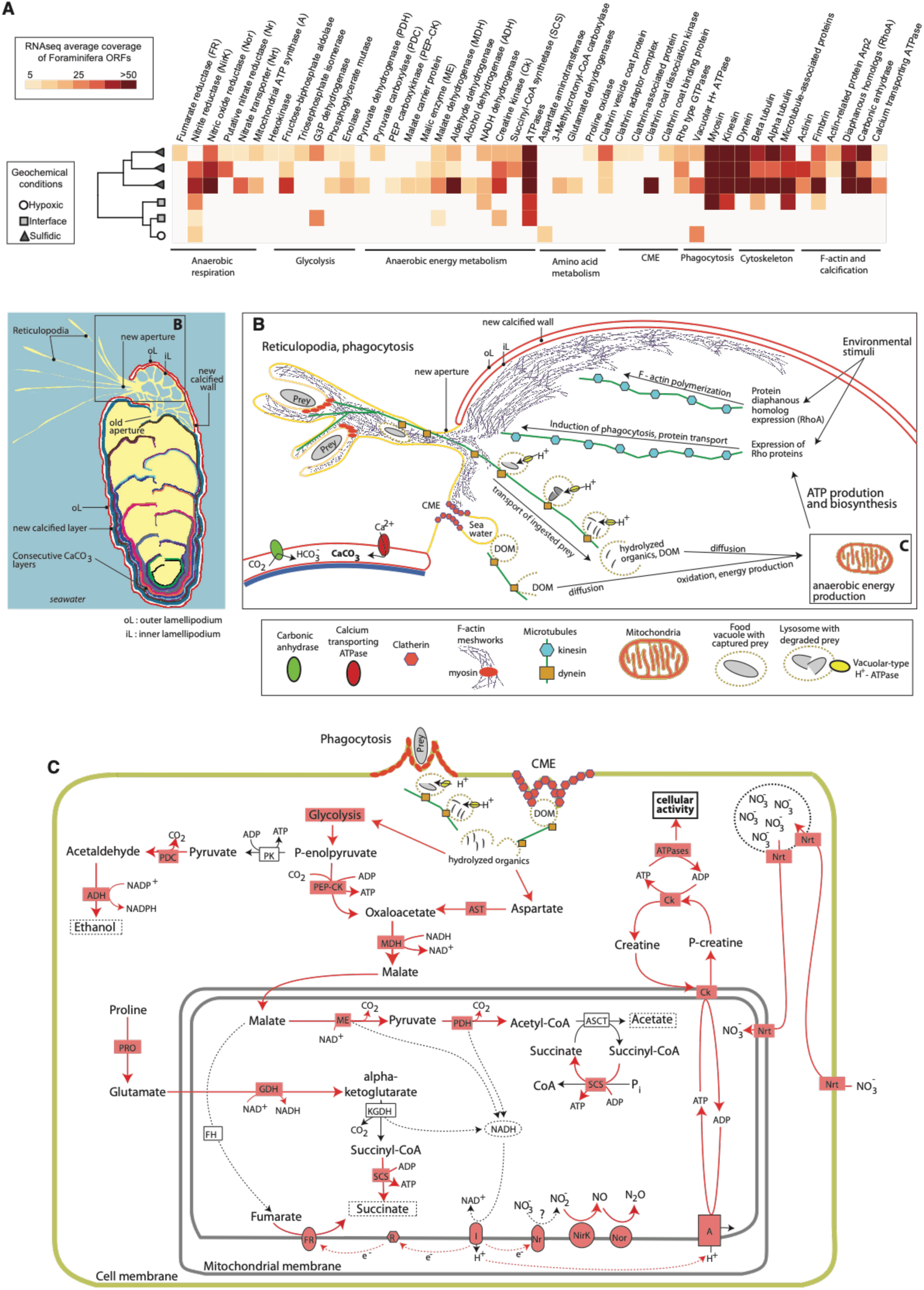
Expression of Foraminifera ORFs involved in key anaerobic physiologies. (A) Heatmap displaying the expression levels of Foraminifera ORFs involved in anaerobic energy production and physiology. Dendogram shows hierarchical clustering (UPGMA) of the samples based on the RNAseq data. One metatranscriptome from the core top and one from the 12 cm sample did not have any detectable expression of the ORFs of interest and are thus not shown. (B) Reconstruction of anaerobic cellular activities in Foraminifera including biomineralization, phagocytosis, CME, and transport of ingested cargo (e.g., prey cells) based on the gene expression data shown in panel A. (C) Reconstruction of potential anaerobic energy production pathways in Foraminifera based on the gene expression data shown in panel A. Red colors show genes that were expressed, red arrows show reactions that are predicted to occur based on the expression of the corresponding gene. Where expressed, gene abbreviations (e.g., Nrt) are shown in red boxes, that correspond to the same labels in panel A. Gene abbreviations that are not highlighted in red are present in tthe genome of the benthic foraminifera species *Globobulimina turgida* and *G. auriculata* ^11^, but their expression was not detected. These include FH: fumarase, KGDH: alpha-ketoglutarate dehydrogenase, PK: pyruvate kinase, and ASCT: acetate:succinate CoA-transferase. The nitric oxide reductase (Nor) gene is not encoded in the benthic Foraminifera genome, but its true absence is uncertain ^11^. This updated representation of Foraminifera anaerobic energy production is modified from anaerobic energy metabolism pathways in eukaryotes that were previously reviewed ^39,40^.

The Foraminifera gene expression data indicate four possible anaerobic mechanisms of ATP production in benthic Foraminifera: [1] substrate level phosphorylation (SLP) of sugars and amino acids via glycolysis and fermentation, [2] dephosphorylation of creatine phosphate via creatine kinase, [3] use of fumarate as a terminal electron acceptor via fumarate-NADH reductase, and [4] dissimilatory reduction of nitrite to generate proton gradient at the membrane for generation of ATP via ATP synthase (Fig 4c). A partial foraminiferal denitrification pathway (*10*) was expressed including a putative dissimilatory nitrate reductase (Nr), dissimilatory nitrite reductase (*16*), and nitric oxide reductase (Nos) (Fig 4a). Additionally, genes encoding foraminiferal nitrate transporters (*10*) (Nrt) were expressed indicating active transmembrane nitrate transport (Fig 4). No homologs to NarK type nitrate/nitrite anitporters that common in denitrifying bacteria (*17*), were detected in the Formainifera transcriptomes. Apparently, these anaerobic energy production mechanisms produce sufficient ATP in the Foraminifera cells to fuel energetically costly biosynthesis pathways including production of reticulopodia, phagocytosis, and clathrin mediated endocytosis (Fig. 4).

The anaerobic energy production mechanisms also produce sufficient ATP in the Foraminifera cells to fuel biomineralization (Fig 4). Of note are the expression of Foraminifera ORFs encoding F-actin proteins, that have been shown experimentally to be involved in the biomineralization of the calcium carbonate test (*18*). Foraminiferal genes encoding ORFs with similarity to protein diaphanous homolog 1 (DIAPH1) were also expressed (Fig 4a), which respond to environmental stimuli and are responsible for actin nucleation and elongation factor required for the assembly of F-actin structures (*19*). Since F-actin is required for biomineralization and calcification of the Foraminifera test (*18*), the expression of DIAPH1 is indicative of ongoing calcification in Foraminifera under anoxic conditions. This is consistent with prior experimental evidence that Foraminifera can calcify under anoxia (*20*).

Foraminiferal genes encoding Rho proteins were expressed, that are responsible for the induction of phagocytosis (*21, 22*). Furthermore, Foraminiferal vacuolar-type H+ ATPases were expressed (Fig 4), which are responsible for lysing digested prey cells inside food vacuoles after phagocytosis (*23*) (Fig 4). Foraminifera ORFs were also expressed that encoded microtubules, kinesin, and dynein, the latter two which are responsible for sending and receiving cellular cargo to and from the membrane, respectively (Fig 4). The expression of ORFs encoding “unconventional” (non muscle) myosin I, II, and VII (Fig 4) from Foraminifera further indicate active phagocytosis. These nonmuscle myosins accumulate at the “phagocytic synapse” (Fig 4b), the point of contact between the pseudopodia and prey cell, which suggests a role for contractile motors proteins during particle internalization (*24*). Pseudopod extension and engulfment has been shown experimentally to be mediated by myosin II that is recruited to the phagocytic synapse (*25*). However, in addition to phagocytosis, myosin motor proteins play an important part in several cytoskeletal processes involving movement such as cell adhesion, cell migration and cell division (*26*). Thus, it is likely that myosins expressed by the Foraminifera under anoxic conditions play a role in a wide range of cellular processes that require force and translocation, for example their motility through the sediment matrix as they search for prey. Clathrin-encoding genes from Foraminifera were also expressed in two samples (at 28 cmbsf) that are involved involved in clathrin-mediated-endocytosis (CME), an additional form of endocytosis and involves an invagination of the membrane via clathrin proteins (*23*). CME results in much smaller vesicles (30-200 nm) compared to those obtained from phagocytosis (500 – 9,000 nm) (*23*) and are used to ingest signaling molecules and other forms of dissolved organic matter. Collectively, these data highlight the key cellular processes needed for survival under anoxia in benthic Foraminifera.

A 10-day incubation of sediment collected from the seafloor, showed that benthic Foraminifera increased their gene expression 27 (+/- 9) fold after the development of anoxic conditions within 20 hrs (Fig. 5). This dramatic increase was observed after oxygen consumption declined steadily over the first 20 hours of the incubation, which was consistent between all biological replicates (Fig. 5). After the development of anoxic conditions, Foraminifera gene expression decreased progressively but still remained 10 to 20 times higher than the t_0_ values up for at least 6 days (Fig 5). After 10 days, the gene expression levels decreased further down to 0.36% (+/- 0.07) of total transcripts, but this was still elevated 2-fold relative to the t_0_ values.

**Figure 5.**
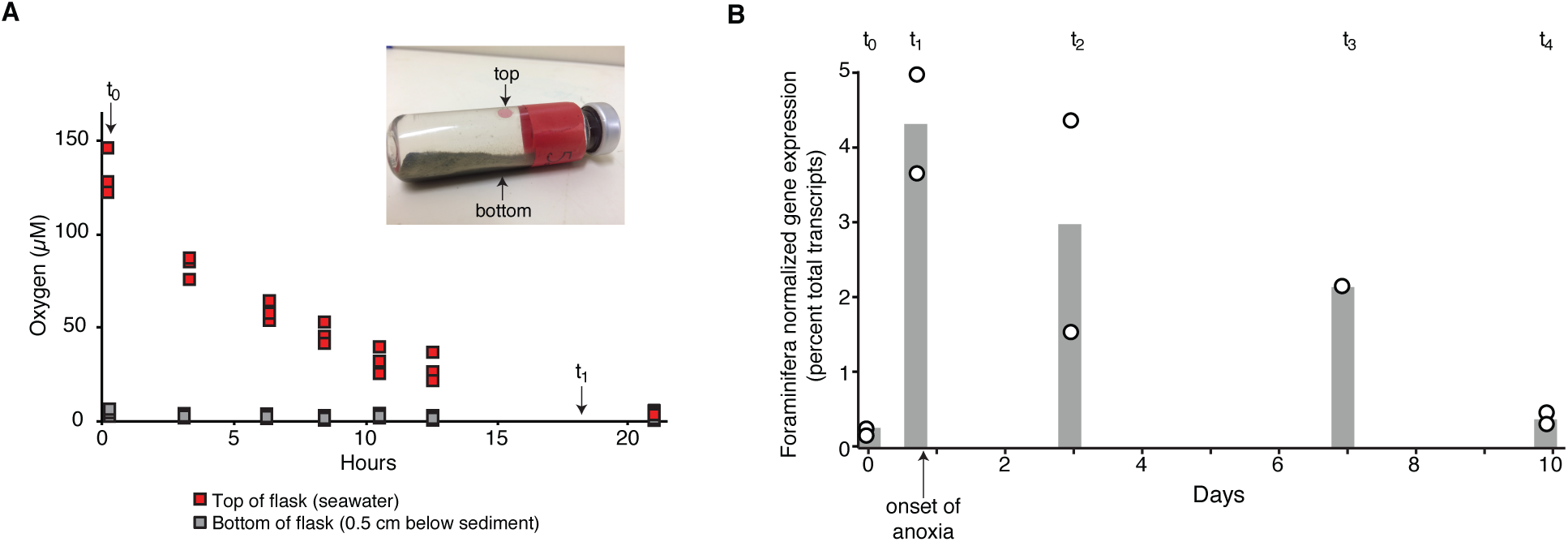
Oxygen consumption and Foraminifera gene expression in a 10-day incubation. (A) Oxygen consumption at the top (in seawater) and bottom (underneath the sediment) of the incubated sediments, the photo shows the experimental setup and the positioning of the two oxygen sensor spots where measurements were made. After the onset of anoxia after 20 hours, the top and bottom of the flask remained anoxic for the duration of the incubation. The flask was incubated in the dark at 10 °C. The individual data points represent O_2_ measurements made on the four separate flasks incubated for the t_1_, t_2_, t_3_, t_4_ timepoints. The 21 hr point includes only the t_2_, t_3_, t_4_ flasks since t_1_ was already taken at 18 hrs. (B) The relative abundance of Foraminifera transcripts (percent of total transcripts) at t_0_ and the four timepoints, individual points represent replicates and histograms represent the average values. Note the sharp increase in gene expression that coincides with the onset of anoxia after 20 hrs.

## Discussion

On the Namibian shelf, Foraminifera live deep below the seafloor down to ca. 30 cmbsf (*27*), co-existing with sulfate reducing bacteria in an anoxic environment that is extremely high in sulfide (*28*). The steadily decreasing abundance of Foraminifera cells in the core with anoxic conditions (Fig 1) is consistent with the reduced rate of heterotrophic metabolism in Foraminifera under anoxic conditions (*29*), and lower levels of ATP in many Foraminifera under anoxia (*11*). The dominance of *Bolivina* throughout the core and our detection of their 18S rRNA, even into the anoxic depths, is consistent with the known affinity of *Bolivina* for oxygen-depleted habitats (*30*), including the studied region as it was observed previously in multiple coring locations on the Namibian shelf (*27*). The “trophic oxygen model” developed by Jorissen et al (*31*) predicts that the dynamic nature of microhabitats allows Foraminifera to migrate up and down in the sediment with the prevailing redox conditions, which is controlled by the organic matter flux (*32, 33*). Hence, since we sampled during the southern Winter when bottom water oxygen levels in the Namibian OMZ are higher (*34, 35*), it is possible that the penetration depth of the Foraminifera extends relatively deep because of the higher oxygen concentration at the sediment surface.

Although the diversity of Foraminifera is well constrained by morphological studies, the group is not yet well represented in transcriptomic and genomic databases. The recently large transcriptome sequencing effort of microbial eukaryotes helped to alleviate this problem (*14*), since it included several Foraminifera that we could add to our database. Nevertheless, because of the relatively low number of sequenced genomes and transcriptomes from Foraminifera (compared to bacteria for example), our metatranscriptome approach cannot distinguish between ORFs derived from different Foraminifera species. The ORFs assigned to Foraminifera here thus serves as a “group averaging”, but should correspond to genetically similar populations since the *de novo* assemblies that are used to build the contigs from the RNAseq data are based on genetic similarity (see Methods). Furthermore, our metatranscriptomes contained the complete 18S rRNA sequence (Fig. 3) from the most abundant taxa, i.e., *Bolivina* sp. and *Stainforthia* sp. (Fig S2) and thus we are confident that the ORFs assigned as Foraminifera are derived primarily from these cytoplasm-containing Foraminifera tests that we could enumerate in the core (Fig. 1). It is further worth mentioning that despite the presence of two morphological different *Bolivina* species in the core, we could not find signs for the active expression of the 18S rDNA in the second species. This indicates that most of the identified foraminiferan metatranscriptomic expression likely comes from one of the *Bolivina* species in addition to *Stainforthia* sp. These findings implies that one foraminiferan species can be active under anoxic conditions while a congeneric species might not be (as) active.

Foraminifera are predators, and are thought to act primarily as heterotrophs utilizing ingested prey cells as carbon sources for growth (*36*). Our gene expression analysis provides insights into the mechanisms of prey acquisition, and the metabolic processing of the ingested material. The expression of ORFs encoding Rho proteins by Foraminifera indicate an active induction of phagocytosis, since Rho proteins function in actin dynamics during phagocytosis (*21, 22*). Myosin motor proteins are recruited to the cell membrane during phagocytosis in order to envelope and capture prey particles (*37*), and the prey then enter the phagocytosing cell as a food vacuole (*23*). Food vacuoles are then transported in to the cell via dynein along microtubules, where the contents are digested under acidic conditions via the activity of vacuolar-type H^+^ ATPases (*23*) (Fig. 4). Such proton pumping ATPases are responsible for lysing digested prey cells inside food vacuoles after phagocytosis, where the acidified lysosomal vesicles are loaded with digestive enzymes (*23*). The metatranscriptome data indicate that under anoxic conditions, the Foraminifera metabolize the hydrolyzed organics for ATP production via fermentation and fumarate reduction, and dissimilatory nitrite reduction (Fig. 4). Because cells are mostly protein, anaerobic fermentation of ingested prey cells by Foraminifera may include amino acid fermentations. By weight, exponentially growing cells are made of roughly 50-60% protein, 20% RNA, 10% lipids, 3% DNA, 10-20% sugars as cell wall constituents, and some metabolites (*38-40*). Amino acid fermentations provide roughly one net ATP per amino acid fermented (*23*).

In addition to hydrolyzed organics from ingested prey, the transcriptomes suggest that CME is another mechanism by which Foraminifera could utilize both high- and low-molecular weight dissolved organic matter (dissolved in the pore water of the sediments) under anoxic conditions. Experiments using ^13^C-labeled diatom prey showed that under anoxic conditions the benthic foram *Ammonia tepida* reduced the number of phagocytosed diatom cells, and the ingested cells were apparently not digested inside vacuoles but remained intact after 4 weeks (*29*). If a decreased utilization of ingested prey for energy production is a general feature of anaerobic Foraminifera, it is possible that organic matter obtained via CME (Fig. 4b, c) becomes a relatively more important carbon source as opposed to ingested prey cells.

Eukaryotic fermentations can produce a variety of end products, and our data indicate the possibility for Foraminifera to produce ethanol, acetate, succinate (Fig. 4c). Under conditions of prolonged anaerobiosis, propionate is preferentially formed as opposed to succinate in anaerobic mitochondria, whereby one additional ATP and one CO_2_ are formed from D-methylmalonyl-CoA via propionyl-CoA carboxylase (*41, 42*). We detected expression of a Foraminifera ORF with similarity to propionyl-CoA carboxylase at 28 cmbsf (data not shown) indicating that prolonged anoxic conditions stimulate production of propionate in Foraminifera mitochondria.

A key intermediate in the anaerobic energy metabolism of most eukaryotes is malate (*41, 42*). During anaerobic respiration in many eukaryotes malate is converted to fumarate via the enzyme fumarase running in reverse, and the resulting fumarate then can be used as the terminal electron acceptor (*41, 42*). This fumarate reduction is coupled to an anaerobic electron transport chain in which electrons are transferred from NADH to fumarate via a specialized complex I and a mitochondrial membrane associated fumarate reductase (*41, 42*). This physiology is typical of anaerobic mitochondria, that exist in the Foraminifera species *Valvulineria* and *Gromia*, and are widely distributed amongst eukaryotes including Bivalvia, Polychaeta, Platyhelminthes, Nematoda, Euglenida, and Ciliophora (*41*).

The metatranscriptomes furthermore indicated that under anoxic conditions, Foraminifera utilize creatine kinase and phosphocreatine to maintain cellular energy homeostasis (Fig. 4c). In eukaryotic cells, creatine kinase acts as a mechanism for maintaining balance between ATP consuming and producing processes (*43*). Our data indicate that this also occurs in anaerobic Foraminifera. In human cells, creatine kinase acts as an ATP regenerator, and the phosphocreatine pool is used as a temporal energy buffer to maintain ATP/ADP ratios inside the cell (*43*). By acting as an energy shuttle between ATP providing and consuming processes, phosphocreatine might help facilitate more energetically costly cellular activities under anoxic conditions for the Foraminifera, such as phagocytosis, by maintaining the spatial “energy circuit” (*44*). For example, creatine kinase contributes to the build-up of a large intracellular pool of phosphocreatine that represents an efficient temporal energy buffer and prevents a rapid fall in global ATP concentrations (*43*). This likely helps to couple the energy producing and energy consuming processes inside of Foraminifera cells during anaerobic metabolism.

Biogeochemical studies indicate that foraminiferans are capable of performing denitrification, that is, the conversion of NO_3_^−^ to N_2_ (*9*). The enzymes behind the foraminiferal denitrification pathway in the genus *Globobulimina* appear to be acquired relatively early in Foraminifera evolution (*10*), and it was indicated that the foraminifera themselves, not associated prokaryotes, are performing the denitrification reaction (*45*). The sequestration of nitrate by Foraminifera is highly suggestive that the protists themselves, and not associated symbionts, are performing nitrate respiration (*45*).

Consistent with this prior evidence, we found the genes of the denitrification pathway in Foraminifera to be expressed (Fig. 4). One of these genes was shown to be a putative assimilatory nitrate reductase (Nr), but it may however function as a sulfite oxidase or dissimilatory nitrate reductase (*10*). Evidence for the potential dissimilatory nitrate reduction comes from this enzyme being shown to catalyze denitrification in the fungus *Cylindrocarpon tonkinense* under specific conditions (*46*). As described previously, we interpret the Nr genes to be involved in dissimilatory nitrate reduction with caution and refer to them as “putative nitrate reductases” since it is possible that the Nr genes function solely for nitrate assimilation in Foraminifera (*10*). In any case, our data show that these Nr genes are transcribed during anaerobic metabolism in benthic Foraminifera.

The expression of nitrate transporters (Nrt) from Foraminifera at 28 cmbsf (Fig. 4a) seems contradictory to the geochemical conditions, since nitrate and nitrite were both below detection at this depth in the core (Fig 1). However, this can be explained by the fact that many benthic Foraminifera can store nitrate in vacuoles under anoxic conditions and use the stored nitrate and nitrite as terminal electron acceptors for anaerobic respiration (*9, 45, 47*). Thus, the expression of the nitrate transporter genes seen here could be responsible for transporting nitrate out of the vacuole (and regulating the cytosolic concentration of nitrate), and into the mitochondrion, as has been proposed previously for denitrifying Foraminifera based on genome data (*10*). The expression of the NirK and Nor genes indicate that the Foraminifera were actively performing two key steps of denitrification – nitrite and nitric oxide reduction (Fig 4c). Some *Bolivina* and *Stainforthia* and species lack a nitrous oxide reductase and reduce nitrate only to N_2_O (*45, 48*), and we did not detect any expression of NosZ indicating that the denitrifying *Bolivina* and *Stainforthia* species in our samples were also likely reducing nitrite to nitric oxide, that is then reduced to N_2_O via Nor (Fig 4c). The lack of expression of the NosZ gene raises the possibility that the denitrifying Foraminifera in Namibian sediments are a source of N_2_O, an important greenhouse gas (*49, 50*). This might be a common feature of denitrifying eukaryotes in the benthos, since denitrifying Fungi in marine sediments also do not contain a nitrous oxide reductase and are an important source of N_2_O (*51*).

The large increase in Foraminifera gene expression upon the onset of anoxic conditions in the incubation (Fig 5) provides experimental support for the observation of increasing Foraminifera gene expression with increasing depths and sulfidic conditions in the core (Fig 2). Thus, the transcriptional activity of many benthic Foraminifera is indeed stimulated by anoxic conditions, which is consistent with experiments that showed benthic Foraminifera can survive for at least 80 days under anoxic conditions with H_2_S (*8, 47*). The peak stimulation of Foraminifera gene expression after 18 hrs at the onset of anoxic conditions might indicate the utilization of nitrate and or nitrite by anaerobic denitrifying foraminifera, once the oxygen had been consumed to below detection values. This indicates that the *Bolivina* and *Stainforthia* species in the Namibian sediments are anaerobes that prefer anoxic conditions, as this clearly stimulated their activity compared to aerobic conditions.

## Conclusions

The increased gene expression by Foraminifera under sulfidic conditions shows for the first time that some foraminifera apparently not only survive, but are thriving, under anoxic conditions in the seafloor. Looking at the data, it becomes evident that the anaerobic energy metabolism of these Foraminifera is sufficient to support phagocytosis, clathrin-mediated-endocytosis, and biocalcification under anoxia. The data also confirm that clades of *Stainforthia* and *Bolivina* utilize pathway for denitrification and identified four pathways of ATP generation including [1] substrate level phosphorylation and fermentation, [2] fumarate reduction, [3] dissimilatory nitrate reduction, and [4] dephosphorylation of creatine-phosphate. This all indicates that anoxic sediments are a primary habitat of some benthic Foraminifera where they are capable to perform all necessary cellular functions. This anaerobic metabolism is consistent with the evidence for the emergence of Rhizaria in the Precambrian where widespread oxygen depletion was present (*52*). This aided the survival of benthic Foraminifera over multiple mass extinctions over the last 500 million years associated with oxygen depletion, thus enabling the utility of their preserved tests as important proxies for paleoclimate and paleoceanography.

## Methods

### Sampling

A 30 cm long sediment core was obtained from a water depth of 125 m the Namibian continental shelf (18.0 S, 11.3 E) during *F/S* Meteor Expedition M148-2 ‘EreBUS’ on July 10^th^, 2018. In brief, the core was acquired with a multi corer (diameter 10 cm), which yielded an intact sediment/water interface and the upper 30 cm of sediment. After retrieval, cores were moved immediately to a 4 °C cold room and sliced every 2 cm within 24 hours. Sections were transferred immediately into sterile, DNA/RNA free 50 mL falcon tubes and then frozen immediately at -20 °C until DNA and RNA extractions. Pore water geochemistry measurements were performed acquired from the same core, methodology and data have been published elsewhere (*12*) and the results are reported in this publication in the Figure 1B.

### Cell counting and enumeration

Between 1 and 4 grams of deep-frozen sediment from 9 sediment depths were thawed and washed over a 63 micron mesh. The residue was immediately wet-sorted and tests of cytoplasm containing Foraminifera were separated, identified to a genus level following Altenbach and Leiter (2010) and enumerated. Representative specimens were photographed using a KEYENCE VHX-6000.

### RNA extraction

RNA was extracted as previously described (*12*). In brief, RNA was extracted from 0.5 g of sediment using the FastRNA Pro Soil-Direct Kit (MP Biomedicals) following the manufacturer’s instructions with final elution of templates in 40 µL PCR water (Roche) as described previously (*12*) with some modifications to maximize RNA yield and reduce DNA contamination. The first modification was that, after the supernatant was removed after first homogenization step, a second homogenization was performed with an additional 500 µL RNA Lysing Buffer. The tubes were centrifuged once again for 5 minutes at maximum speed, and the supernatant from the second homogenization was combined with that resulting from the first homogenization, continuing with the protocol from the manufacturer. Second, we added glycogen at a concentration of 1 µg/mL during the 30-minute isopropanol precipitation in order to maximize recovery of the RNA pellet. To reduce DNA contamination, we extracted all RNA samples in a HEPA-filtered laminar flow hood dedicated only for RNA work (no DNA allowed inside) that also contains dedicated RNA pipettors used exclusively inside the hood with RNA samples. All surfaces were treated with RNAse-Zap prior to extractions and exposed to UV light for 30 minutes before and after each extraction.

### Metatranscriptomics

Metatranscriptomes were prepared as previously described (*12*). In brief, DNAse treatment, synthesis of complementary DNA and library construction were obtained from 10 µL of RNA templates by processing the Trio RNA-Seq kit protocol (NuGEN Technologies). Libraries were quantified on an Agilent 2100 Bioanalyzer System, using the High Sensitivity DNA reagents and DNA chips (Agilent Genomics). The libraries constructed using specific (different) barcodes, pooled at 1 nM, and sequenced in two separate sequencing runs with a paired-end 300 mid output kit on the Illumina MiniSeq. A total of 40 million sequences were obtained after Illumina sequencing, which could be assembled *de novo* into 41,230 contigs. Quality control, *de novo* assembly, and ORFs searches were performed as described previously (*12*), with the additional step of using the eukaryotic code for translations and ORF predictions.

### Gene identification

A total of 8,556 ORFs were found that were then searched for similarity using BLASTp against a database (*12*) containing predicted proteins from all protist, fungal, bacterial, and archaeal genomes and MAGs in the JGI and NCBI databases using DIAMOND (*53*). This database also contained all ORFs from the >700 transcriptomes of microbial eukaryotes from the MMETS project (*14*) and the recently published foraminiferal genome and transcriptome containing the novel denitrification pathway (*10*). Cutoff for assigning hits to specific taxa were a minimum bit score of 50, minimum amino acid similarity of 30, and an alignment length of 50 residues. Extraction blanks were also sequenced alongside the environmental samples to identify contamination, and ORFs from contaminant taxa. We assigned ORFs as being derived from Foraminifera if they had a significant similarity above this threshold to a predicted protein from a previously sequenced Foraminifera transcriptome or genome. Because our database contains predicted proteins from >700 transcriptomes of other microbial eukaryotes, we are confident that this level of stringency is sufficient to make a broad level of taxonomic assignment of ORFs from the metatranscriptomes to Foraminifera in general (as opposed to being actually derived from other protist groups).

ORFs assigned as Foraminifera were then additionally annotated against the Cluster of Eukaryotic Orthologous Genes (KOG) database (*15*), using DIAMOND with the same parameters as above. The lack of metatranscriptomic ORFs having highest similarity to *Bolivina* and *Stainforthia* (Fig S1) is easily explained by the lack of transcriptome data from the species in public databases. Nevertheless, because we cannot be sure from which species each of our metatranscriptome ORF derives, we annotated all of the ORFs having highest similarity to a previously sequenced Foraminifera transcriptome or genome, as being derived from Foraminifera.

Contamination in the metatranscriptomes were primarily diatoms (“lab weeds”), cyanobacteria, *Streptococcus, Acinetobacter, Staphylococcus, Rhizobium, Ralstonia*, and *Burkholderia*. All ORFs that were shared between contaminant samples and the metatranscriptomes were removed prior to analysis. Incorporation of protist transcriptomes^49^ greatly reduced the amount of laboratory contamination from eukaryotic algae such as diatoms (“lab weeds”) introduced during the library prep. All metatranscriptomes had <10% ORFs from contaminating taxa.

### Incubation experiment

Immediately after core retrieval and freezing of the core top samples, 2 g aloquots of sediment from the core top was added to four 20 mL sterile glass vials (for t_1_, t_2_, t_3_, t_4_ timepoints) containing sterile oxygen sensor spots (PreSens Precision Sensing). Oxygen was measured non-invasively using the Fibox (PreSens Precision Sensing) as described previously (*54*). The sediment was overlaid with ca. 18 mL of the natural hypoxic bottom water collected in the multicore leaving no air in the headspace, and crimp sealed with grey rubber butyl stoppers. The flasks were incubated on the side and oxygen sensor spots were positioned at the top (to measure oxygen in the overlying seawater) and bottom (to measure oxygen at the base of the sediment) of the flask (see Fig 5 for a photo of the setup). The flasks were incubated in the dark at 10 °C and taped to the surface of the bench to prevent rolling and mixing of the tube. Each of the four flasks for the timepoints were frozen separately at the respective timepoints t_1_ (18 hrs), t_2_ (3 days), t_3_ (7 days), and t_4_ (10 days) immediately at -20 °C. Because the incubation was set up immediately after core retrieval and freezing the core top samples, the frozen core top samples served as the t_0_ samples for the start of the incubation. RNA extractions, metatranscriptomes, and bioinformatic processing was performed as described above.

### Phylogenetics

To identify the likely active foraminifera taxa in the sediments, we searched for foraminiferan 18S rDNA OTUs present within the metatranscriptomes. We performed BLASTn searches (Discontiguous Megablast, e-value 1E-10). As query we used a small custom made database of complete foraminifera sequences based on Pawlowski *et al*. (*55*) and Holzmann and Pawlowski (*56*). The resulting OTUs were reciprocally blasted against NCBI’s nr database (Discontiguous Megablast, e-value 1E-10). The two OTUs with highest similarity to Foraminifera 18S rDNA were further used for sequence extensions using a greedy approach. For this, 10bp on both ends were trimmed from the putative foraminiferan 18S rRNA OTUs to remove possible erroneous bases due to dropping read quality towards the ends of reads. We only extended the OTU fragment matching the last 1000bp of the foraminiferan 18S rRNA sequences since this is a commonly used foraminifera barcoding region and allows the comparison with a wide diversity of previously barcoded foraminiferan taxa (*57*). We performed 20 iterations of greedy extension in GENEIOUS Prime 2019 (*58*) by mapping trimmed metatranscriptomics reads (trimmed with TRIMMOMATIC v.0.38 (Bolger, 2014 #5074) and default options) to the end-trimmed 18S rDNA OTUs. This extended 5’ and 3’ ends of the 18S rRNA OTUs. Both sequences were manually error corrected based on the mapped reads. We carefully and manually proved that read pairs spanned regions of high sequence similarity with other foraminiferans, i.e. highly conserved stem regions of the 18S rRNA. This approach allowed us to unambiguously extend both OTUs to yield the full 18S rRNA barcoding region. These sequences were blasted against the NCBI nr database and showed strong sequene similarity to the benthic foraminifera genera *Stainforthia* and *Bolivina*. In order to confirm their taxonomic affiliation and to refine their placement, we established two separate alignment that included 30 sequences of the genus *Bolivina* (*59*) on the one hand, and on the other hand 30 sequences of sister genus *Stainforthia* (*56*). The two separate sequence sets were automatically aligned with MAFFT v.7 (*60*) and a phylogenetic inference was calculated with 1000 non-parametric bootstrapping pseudo replicates based on a BioNJ starting tree using PhyML (*61*). The best substitution models were automatically selected using the Smart Model Selection (*62*) under Akaike Information Criterion and the model GTR+I+G was selected for the *Bolivina* alignment and the model TN93+G+I was selected for the *Stainforthia* alignment. Both trees were visualized using ITOL and are provided in Figure 3.

## Acknowledgements

We dedicate this work to the late Prof. Dr. Alexander V. Altenbach, whose legacy of research into anaerobic Foraminifera was a source of inspiration for completing this study. This work was supported by the Deutsche Forschungsgemeinschaft (DFG) through Project OR 417/4-1 (W.D.O), and the *F/S* Meteor Expedition M148/2 ‘EreBUS’. The authors thank the captain and crew of the *F/S* Meteor assistance during the oceanographic expedition, as well as T. Ferdelman, S. Littmann, T. Wilkop, G. Klockgether and K. Imhoff who assisted in obtaining samples. This work was performed in part through the Masters in Geobiology and Paleontology Program (MGAP) at LMU Munich. G.W. acknowledges funding through the LMU Munich’s Institutional Strategy LMUexcellent within the framework of the German Excellence Initiative and the European Union’s Horizon 2020 Marie Skłodowska-Curie Innovative Training Network IGNITE (No. 764840). RM and MK acknowledge funding by the Deutsche Forschungsgemeinschaft (DFG, German Research Foundation) through Germany’s Excellence Strategy (EXC-2077, grant no 390741603).

## Author contributions

W.D.O, conceived the idea for the study and wrote the paper. R.M., A.V., and T. G. F. produced data. W.D.O., R.M., A.V., M.E., G.W., and T. G. F. analyzed data. All authors participated in editing the manuscript and interpreting the results.

## Additional information

Supplementary Information includes supplemental figures Fig S1 and S2 and Table S1. All sequence data is publicly accessible in NCBI through BioProject number PRJNA525353.

## Correspondence and requests for materials

should be addressed to W.D.O.

## Figures

**Figure S1:**
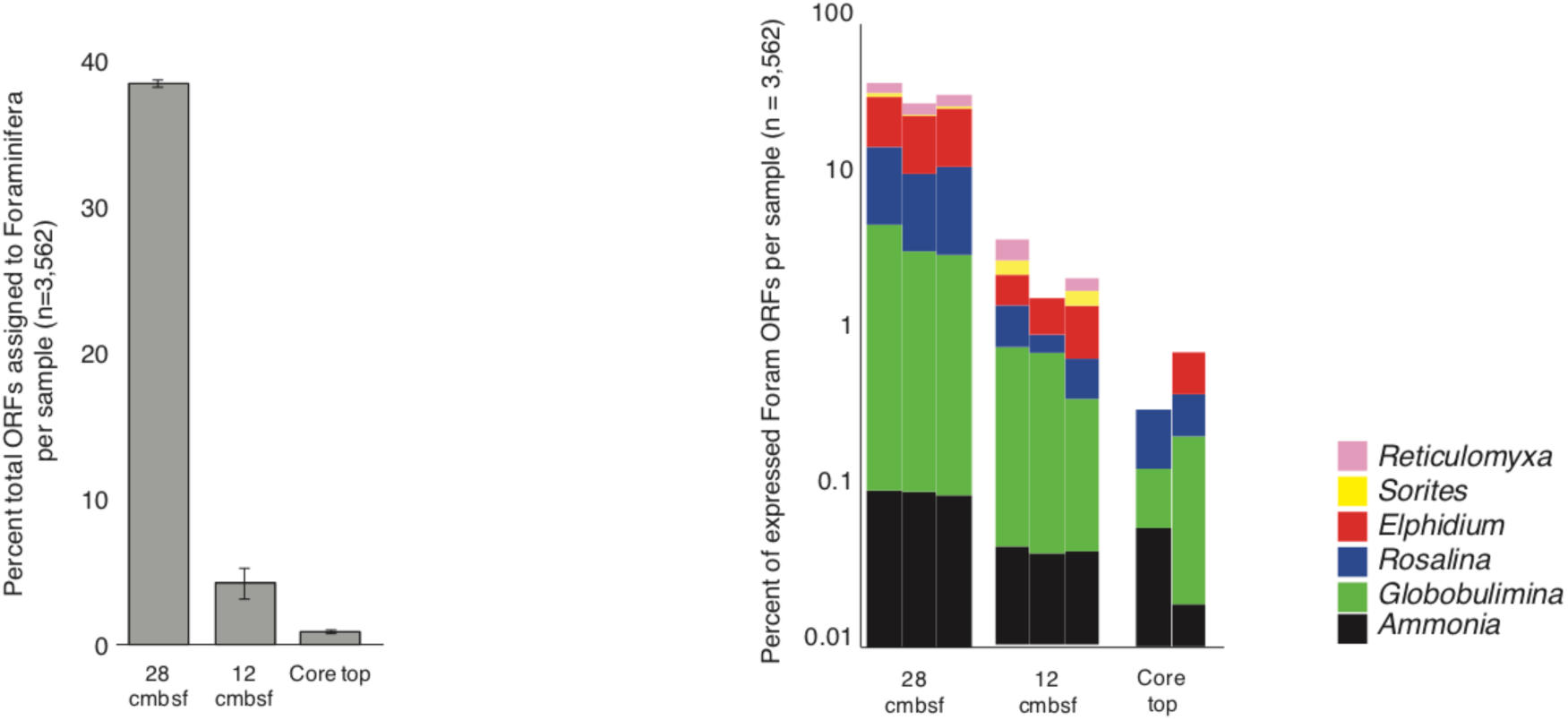
Relative distributions of taxa that Foraminifera-derived ORFs in the metatranscriptome had as top hits after searches with DIAMOND.

**Figure S2.**
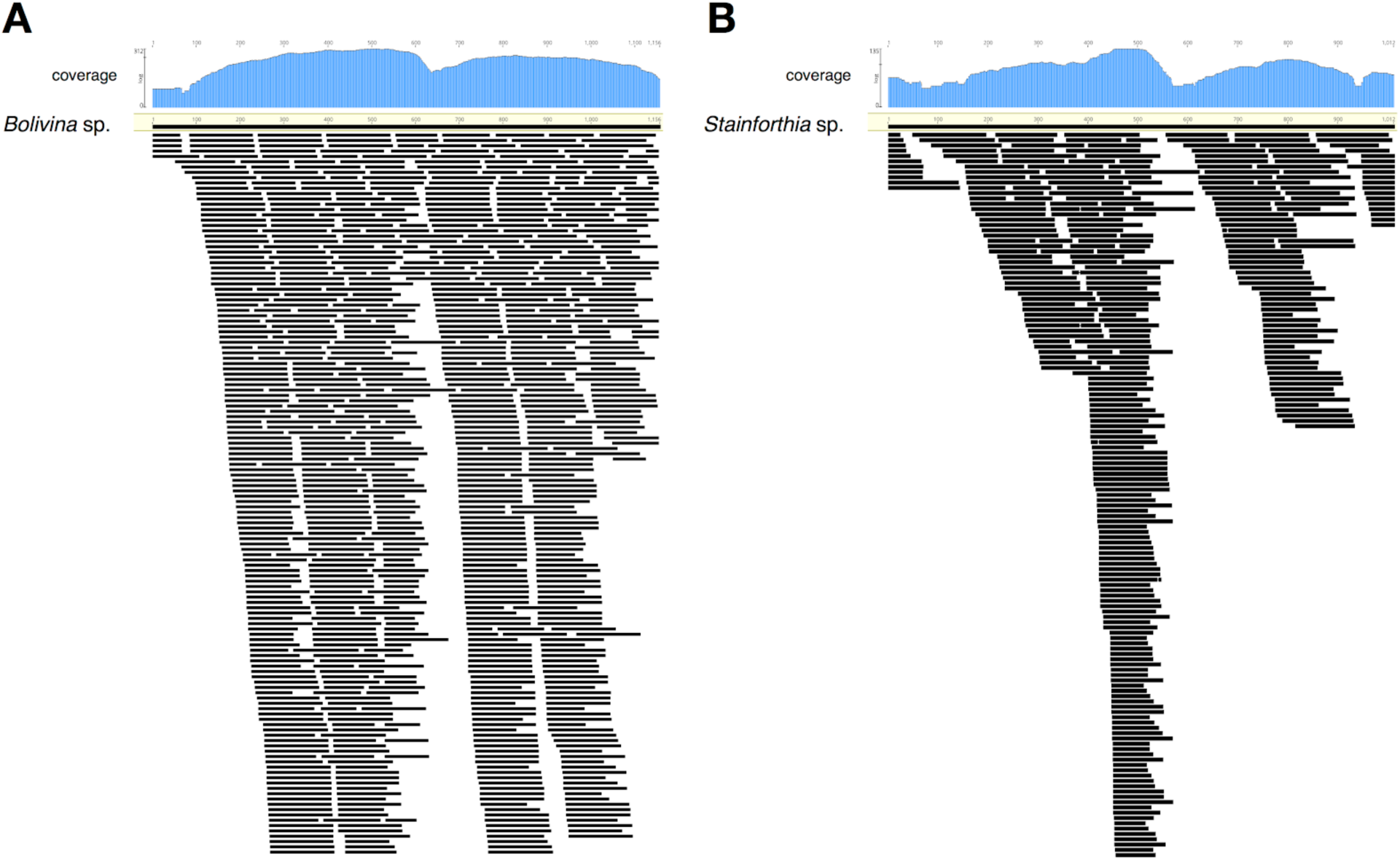
Schematic representation of trimmed metatranscriptomic reads mapped to the previously sequenced 18S rRNA genes *Bolivina* sp. (A) and *Stainforthia* sp. (B). The coverage of each fragment is indicated with blue histograms on the top. Each read is shown as black bar. Mapping was performed with GENEIOUS prime as indicated in the methods section. Note that more reads map to Bolivina, which is consistent with the dominance of cytoplasm bearing tests from Bolivina throughout the core (Fig 1).

**Table S1.**
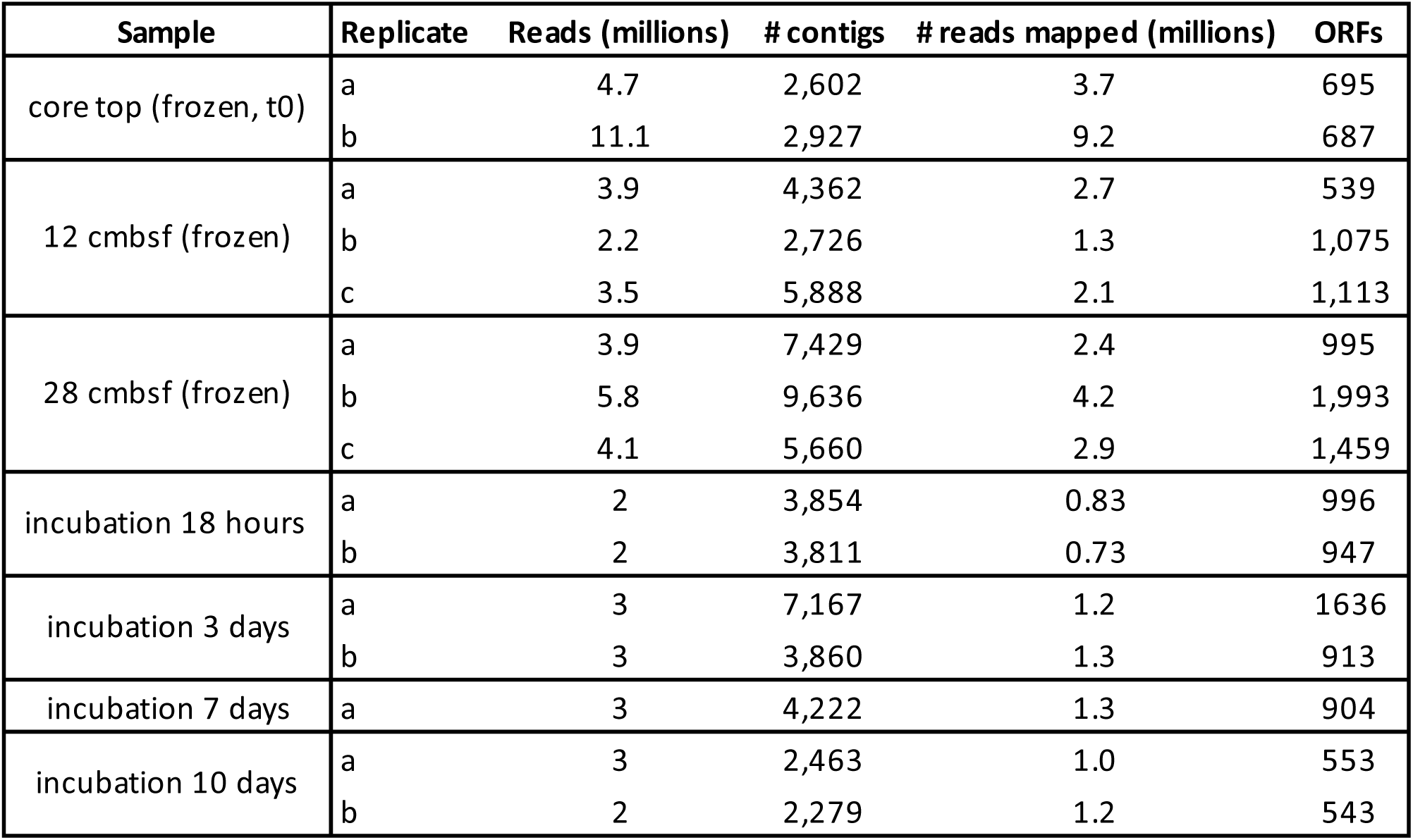
Sequencing and assembly statistics.

| Sample | Replicate | Reads (millions) | # contigs | # reads mapped (millions) | ORFs |
| --- | --- | --- | --- | --- | --- |
| core top (frozen, t0) | a | 4.7 | 2,602 | 3.7 | 695 |
|  | b | 11.1 | 2,927 | 9.2 | 687 |
| 12 cmbsf (frozen) | a | 3.9 | 4,362 | 2.7 | 539 |
|  | b | 2.2 | 2,726 | 1.3 | 1,075 |
|  | c | 3.5 | 5,888 | 2.1 | 1,113 |
| 28 cmbsf (frozen) | a | 3.9 | 7,429 | 2.4 | 995 |
|  | b | 5.8 | 9,636 | 4.2 | 1,993 |
|  | c | 4.1 | 5,660 | 2.9 | 1,459 |
| incubation 18 hours | a | 2 | 3,854 | 0.83 | 996 |
|  | b | 2 | 3,811 | 0.73 | 947 |
| incubation 3 days | a | 3 | 7,167 | 1.2 | 1636 |
|  | b | 3 | 3,860 | 1.3 | 913 |
| incubation 7 days | a | 3 | 4,222 | 1.3 | 904 |
| incubation 10 days | a | 3 | 2,463 | 1.0 | 553 |
|  | b | 2 | 2,279 | 1.2 | 543 |

## References

1. A. V. Altenbach, in Encyclopedia of Geobiology, J. Reitner, V. Thiel, Eds. (Springer, Netherlands, 2012), pp. 393–396.

2. F. Wiese, J. Reitner, in Encyclopedia of Geobiology, J. Reitner, V. Thiel, Eds. (Springer, Dordrecht, Netherlands, 2012), pp. 293–306.

3. J. W. Murray, Ecology and applications of benthic foraminifera. (Cambridge University Press, 2006).

4. A. J. Gooday, L. A. Levin, P. Linke, T. Heeger, in Deep-Sea Food Chains and the Global Carbon Cycle, G. T. Rowe, V. Pariente, Eds. (Springer, Netherlands, 1992), pp. 63–91.

5. A. J. Gooday, H. Nomaki, H. Kitazato, Modern deep-sea benthic foraminifera: a brief review of their morphology-based biodiversity and trophic diversity. Geol Soc Lond Spec Publ 303, 97–119 (2008).

6. A. J. Gooday, J. M. Bernhard, L. A. Levin, S. B. Suhr, Foraminifera in the Arabian Sea oxygen minimum zone and other oxygen-deficient settings: taxonomic composition, diversity, and relation to metazoan faunas. Deep Sea Res Part II Top Stud Oceanogr 47, 25–54 (2000).

7. L. A. Levin et al., Effects of natural and human-induced hypoxia on coastal benthods. Biogeosciences 6, 2063–2098 (2009).

8. L. Moodley, G. J. Van der Zwaan, P. M. J. Herman, L. Kempers, P. Van Breugel, Differential response of benthic meiofauna to anoxia with special reference to Foraminifera (Protista: Sarcodina). Marine Ecology Progress Series 158, 151–163 (1997).

9. N. Risgaard-Petersen et al., Evidence for complete denitrification in a benthic foraminifer. Nature 443, 93–96 (2006).

10. C. Woehle et al., A Novel Eukaryotic Denitrification Pathway in Foraminifera. Curr Biol 28, 2536–2543 e2535 (2018).

11. J. M. Bernhard, E. Alve, Survival, ATP pool, and ultrastructural characterization of benthic foraminifera from Drammensfjord (Norway): response to anoxia. Marine Micropaleontology 28, 5–17 (1996).

12. W. D. Orsi et al., Metabolic activity analyses demonstrate that Lokiarchaeon exhibits homoacetogenesis in sulfidic marine sediments. Nat Microbiol, (2020).

13. L. Pillet, J. Pawlowski, Transcriptome analysis of foraminiferan Elphidium margaritaceum questions the role of gene transfer in kleptoplastidy. Mol Biol Evol 30, 66–69 (2013).

14. P. J. Keeling et al., The Marine Microbial Eukaryote Transcriptome Sequencing Project (MMETSP): illuminating the functional diversity of eukaryotic life in the oceans through transcriptome sequencing. PLoS Biol 12, e1001889 (2014).

15. R. L. Tatusov, E. V. Koonin, D. J. Lipman, A genomic perspective on protein families. Science 278, 631–637 (1997).

16. I. Monirith et al., Asia-Pacific mussel watch: monitoring contamination of persistent organochlorine compounds in coastal waters of Asian countries. Mar Pollut Bull 46, 281–300 (2003).

17. W. G. Zumft, Cell biology and molecular basis of denitrification. Microbiol Mol Biol Rev 61, 533–616 (1997).

18. J. Tyszka et al., Form and function of F-actin during biomineralization revealed from live experiments on foraminifera. Proc Natl Acad Sci U S A, (2019).

19. S. Stritt et al., A gain-of-function variant in DIAPH1 causes dominant macrothrombocytopenia and hearing loss. Blood 127, 2903–2914 (2016).

20. M. P. Nardelli et al., Experimental evidence for foraminiferal calcification under anoxia. Biogeosciences 11, 4029–4038 (2014).

21. G. J. Doherty, H. T. McMahon, Mechanisms of endocytosis. Annu Rev Biochem 78, 857–902 (2009).

22. G. Chimini, P. Chavrier, Function of Rho family proteins in actin dynamics during phagocytosis and engulfment. Nat Cell Biol 2, E191–196 (2000).

23. W. F. Martin, A. G. M. Tielens, M. Mentel, S. G. Garg, S. V. Gould, The physiology of phagocytosis in the context of mitochondrial origin. Microbiol Mol Biol Rev 81, e00008–00017 (2017).

24. O. I. Stendahl, J. H. Hartwig, E. A. Brotschi, T. P. Stossel, Distribution of actin-binding protein and myosin in macrophages during spreading and phagocytosis. J Cell Biol 84, 215–224 (1980).

25. R. K. Tsai, D. E. Discher, Inhibition of “self” engulfment through deactivation of myosin-II at the phagocytic synapse between human cells. J Cell Biol 180, 989–1003 (2008).

26. M. Vicente-Manzanares, X. Ma, R. S. Adelstein, A. R. Horwitz, Non-muscle myosin II takes centre stage in cell adhesion and migration. Nat Rev Mol Cell Biol 10, 778–790 (2009).

27. C. Leiter, A. V. Altenbach, Benthic Foraminifera from the diatomaceous mud belt off Namibia: characteristic species for severe anoxia. Palaeontologia Electronica 13.2.11A, (2010).

28. H. D. Schulz et al., Dense populations of a giant sulfur bacterium in Namibian shelf sediments. Science 284, 493–495 (1999).

29. C. LeKieffre et al., Surviving anoxia in marine sediments: The metabolic response of ubiquitous benthic foraminifera (Ammonia tepida). PLoS One 12, e0177604 (2017).

30. K. A. Koho, E. Pina-Ochoa, in Anoxia: Evidence for Eukaryote Survival and Paleontological Strategies, A. V. Altenbach, J. M. Bernhard, J. Seckbach, Eds. (Springer, Dordrecht, 2012),chap. 4, pp. 251–285.

31. F. J. Jorissen, H. C. De Stigter, J. G. V. Widmark, A conceptual model explaining benthic foraminiferal habitats. Marine Micropaleontology 26, 3–15 (1995).

32. W. D. Orsi, Ecology and evolution of seafloor and subseafloor microbial communities. Nat Rev Microbiol 16, 671–683 (2018).

33. B. Jørgensen, Mineralization of organic matter in the sea bed – the role of sulfate reduction. Nature 296, 643–645 (1982).

34. G. Lavik et al., Detoxification of sulphidic African shelf waters by blooming chemolithotrophs. Nature 457, 581–584 (2009).

35. M. M. Kuypers et al., Massive nitrogen loss from the Benguela upwelling system through anaerobic ammonium oxidation. Proc Natl Acad Sci U S A 102, 6478–6483 (2005).

36. D. A. Caron et al., Probing the evolution, ecology and physiology of marine protists using transcriptomics. Nat Rev Microbiol 15, 6–20 (2017).

37. P. Rougerie, V. Miskolci, D. Cox, Generation of membrane structures during phagocytosis and chemotaxis of macrophages: role and regulation of the actin cytoskeleton. Immunol Rev 256, 222–239 (2013).

38. J. W. Lengeler, G. Drews, H. G. Schegel, Biology of the Prokaryotes. (Blackwell Science, 1999).

39. F. C. Neidhardt, J. L. Ingraham, M. Schaecter, Physiology of the Bacterial Cell. (Sinauer Associates, 1990).

40. A. H. Stouthamer, A theoretical study on the amount of ATP required for synthesis of microbial cell material.. Antonie van Leeuwenhoek 39, 545–565 (1973).

41. M. Muller et al., Biochemistry and evolution of anaerobic energy metabolism in eukaryotes. Microbiol Mol Biol Rev 76, 444–495 (2012).

42. V. Zimorski, M. Mentel, A. G. M. Tielens, W. F. Martin, Energy metabolism in anaerobic eukaryotes and Earth’s late oxygenation. Free Radic Biol Med, (2019).

43. U. Schlattner, M. Tokarska-Schlattner, T. Wallimann, Mitochondrial creatine kinase in human health and disease. Biochim Biophys Acta 1762, 164–180 (2006).

44. T. Wallimann, M. Wyss, D. Brdiczka, K. Nicolay, H. M. Eppenberger, Intracellular compartmentation, structure and function of creatine kinase isoenzymes in tissues with high and fluctuating energy demands: the ‘phosphocreatine circuit’ for cellular energy homeostasis. Biochem J 281 (Pt 1), 21–40 (1992).

45. E. Pina-Ochoa et al., Widespread occurrence of nitrate storage and denitrification among Foraminifera and Gromiida. Proc Natl Acad Sci U S A 107, 1148–1153 (2010).

46. T. O. Watsuji, N. Takaya, A. Nakamura, H. Shoun, Denitrification of nitrate by the fungus Cylindrocarpon tonkinense. Biosci. Biotechnol. Biochem. 67, 1115–1120 (2003).

47. E. Pina-Ochoa, K. A. Koho, E. Geslin, N. Risgaard-Petersen, Survival and life strategy of foraminifer, Globobulimina turgida, through nitrate storage and denitrification: laboratory experiments. Marine Ecology Progress Series 417, 39–49 (2010).

48. A. Kamp, S. Hogslund, N. Risgaard-Petersen, P. Stief, Nitrate Storage and Dissimilatory Nitrate Reduction by Eukaryotic Microbes. Front Microbiol 6, 1492 (2015).

49. S. W. A. Naqvi et al., Marine hypoxia/anoxia as a source of CH4 and N2O. Biogeosciences 7, 2159–2190 (2010).

50. R. N. van den Heuvel, M. M. Hefting, N. C. Tan, M. S. Jetten, J. T. Verhoeven, N2O emission hotspots at different spatial scales and governing factors for small scale hotspots. Sci Total Environ 407, 2325–2332 (2009).

51. S. D. Wankel et al., Evidence for fungal and chemodenitrification based N2O flux from nitrogen impacted coastal sediments. Nature Communications 8, 15595 (2017).

52. B. J. Nettersheim et al., Putative sponge biomarkers in unicellular Rhizaria question an early rise of animals. Nat Ecol Evol 3, 577–581 (2019).

53. B. Buchfink, C. Xie, D. H. Huson, Fast and sensitive protein alignment using DIAMOND. Nat Methods 12, 59–60 (2015).

54. A. S. Ortega-Arbulu, M. Pichler, A. Vuillemin, W. D. Orsi, Effects of organic matter and low oxygen on the mycobenthos in a coastal lagoon. Environ Microbiol 21, 374–388 (2019).

55. J. Pawlowski, M. Holzmann, J. Tyszka, New supraordinal classification of Foraminifera: Molecules meet morphology. Marine Micropaleontology 100, 1–10 (2013).

56. M. Holzmann, J. Pawlowski, An updated classification of rotaliid foraminifera based on ribosomal DNA phylogeny. Marine Micropaleontology 132, 18–34 (2017).

57. J. Pawlowski, M. Holzmann, A plea for DNA barcoding of Foraminifera. The Journal of Foraminiferal Research 44, 62–67 (2014).

58. M. Kearse et al., Geneious Basic: an integrated and extendable desktop software platform for the organization and analysis of sequence data. Bioinformatics 28, 1647–1649 (2012).

59. M. Kucera et al., Caught in the act: Anatomy of an ongoing benthic-planktonic transition in a marine protist. Journal of Plankton Research 39, 436–449 (2017).

60. K. Katoh, D. M. Standley, MAFFT multiple sequence alignment software version 7: improvements in performance and usability. Mol Biol Evol 30, 772–780 (2013).

61. S. Guindon et al., New algorithms and methods to estimate maximum-likelihood phylogenies: assessing the performance of PhyML 3.0. Syst Biol 59, 307–321 (2010).

62. V. Lefort, J. E. Longueville, O. Gascuel, SMS: Smart Model Selection in PhyML. Mol Biol Evol 34, 2422–2424 (2017).

